# ELMO1 dependent efferocytosis protects from nephrotoxin induced acute kidney injury

**DOI:** 10.64898/2026.03.24.713994

**Authors:** Blandine Baffert, Michal Cholko, Vikram Sabapathy, Pritika Modhukuru, Isaac Heath, Shuqiu Zheng, Jitendra Gautam, Kevin Schneider, Lily Silverman, Mark Okusa, Rahul Sharma, Sanja Arandjelovic

**Author notes:** These authors contributed equally to this work (Blandine Baffert, Michal Cholko). Correspondence should be addressed to: Sanja Arandjelovic, 1340 Jefferson Park Avenue, Pinn Hall 4043, University of Virginia, Charlottesville, VA 22903.

## Abstract

Acute kidney injury (AKI) is a sudden episode of kidney failure linked to a wide range of health conditions. High mortality in AKI highlights the need to identify new therapeutic approaches. Homeostasis in multicellular organisms is exquisitely regulated by phagocytosis of apoptotic cells, also known as ‘efferocytosis’. Apoptotic cells are frequently observed at sites of inflammation, including in AKI. Engulfment and cell motility protein-1 (ELMO1) is a regulator of the actin cytoskeleton that promotes apoptotic cell removal by phagocytes during efferocytosis. Mutations in the human *ELMO1* gene are linked with diabetic nephropathy and, in animal models of this disease, high ELMO1 levels promote renal dysfunction. However, the role of ELMO1 in AKI was not known. Here, we describe the links between *ELMO1* and kidney pathology and test global and tissue-specific ELMO1-deficient mice in models of AKI. While global loss of *Elmo1* expression did not impact the immediate loss of renal function after ischemia-reperfusion elicited AKI, ELMO1 deficiency resulted in increased tissue injury in AKI caused by cisplatin injection. Cisplatin induced robust renal cell apoptosis that was significantly elevated in mice with the global loss of ELMO1, but not in mice with the macrophage-specific *Elmo1* deletion. Using primary cell culture and immunofluorescence approaches, we highlight the role of ELMO1 in efferocytosis by several renal cell types, suggesting possible additive effects during nephrotoxic injury.

## Introduction

Acute kidney injury (AKI) is characterized by a sudden deterioration in renal function and is associated with poor prognosis and high mortality rates (1–3). The pathogenesis of AKI is multifactorial, frequently arising from mechanisms including sepsis, ischemia, and nephrotoxicity, thereby complicating diagnosis and therapeutic management(4–6). Treatment options for AKI are limited and supportive; renal replacement therapy is the only FDA approved treatment for AKI (7–9).

While our knowledge of disease pathogenesis in human AKI remains incomplete, many insights have been gleaned from animal models. Renal injury induced by ischemia and reperfusion in rodents is one of the most utilized, constituting a temporary cessation of renal blood flow, followed by reperfusion, during which inflammation plays a significant role in tissue damage. Alternatives to ischemia/reperfusion(10) are nephrotoxin-induced models, such as those induced by gentamicin(11), cisplatin(12), or folic acid(13). In nephrotoxic models, the severity of renal damage is directly proportional to the dose of the nephrotoxic agent(14–16).

Cisplatin, a commonly used clinical chemotherapy agent, accumulates in renal tubular epithelial cells (RTEC), leading to programmed cell death and loss of kidney function(12,14,17–20). Both apoptosis(21–23) and necrosis(24,25) have been observed in animal models of AKI and are linked to disease pathophysiology(22,26–28). Apoptosis is a non-inflammatory type of cell death that is intricately linked with phagocytosis of apoptotic cells, also known as ‘efferocytosis’(29–31). Prompt clearance by efferocytosis explains why apoptotic cells are rarely observed in healthy tissues(29). In instances of failed efferocytosis, apoptotic cells can progress to secondary necrosis (a non-apoptotic, inflammatory form of cell death) and release their intracellular contents into the surrounding environment to activate inflammatory pathways(32).

Engulfment and cell motility protein-1 (ELMO1) is a critical regulator of phagocytosis and cell migration(33) that has been demonstrated to regulate inflammatory disease development and severity in animal models(34,35). Human genetic studies have identified *ELMO1* as a risk gene for development of diabetic nephropathy, with multiple single nucleotide polymorphisms (SNP) in the *ELMO1* gene linked with disease across diverse populations(36–39). However, ELMO1 role in kidney injury is currently unclear.

In this study, we tested whether ELMO1 contributes to acute kidney injury (AKI) using mice with *Elmo1* deletion in two different models: ischemia/reperfusion injury (IRI) and cisplatin-induced nephrotoxicity (cisplatin-AKI). Surprisingly, we discovered a bifunctional role for ELMO1. In IRI, we found that loss of ELMO1 dampened the inflammatory response but did not alleviate the loss of kidney function. On the other hand, early nephrotoxic damage induced by cisplatin injection was elevated in ELMO1 deficient animals, with increased numbers of apoptotic cells in the kidneys. Through analysis of mice with tissue-specific deletion of *Elmo1*, combined with primary cell culture and histopathological analyses, we identify the role of ELMO1 in efferocytosis across multiple cell types that collectively could contribute to protection from nephrotoxic renal injury.

## Results

### ELMO1 is linked with kidney disease in humans and mice

We first explored the links between the human *ELMO1* gene and kidney disease through analysis of publicly available databases. In addition to the known links between *ELMO1* single nucleotide polymorphisms (SNPs) and diabetic nephropathy(36,37,39), we found *ELMO1* genetic associations with multiple renal abnormalities and kidney disease (**Fig. 1A**). Analysis of the human kidney single cell/nucleus datasets(40) showed that *ELMO1* expression can be observed in multiple renal cell types at baseline, with dynamic changes observed in acute kidney injury (AKI) and chronic kidney disease (CKD) (**Fig. 1B**). We also addressed *Elmo1* expression in kidney disease in mice by analysis of the published dataset from a longitudinal ischemia/reperfusion induced acute kidney injury(41). In the healthy mouse kidney, *Elmo1* expression is high in macrophages, podocytes, endothelial cells, and T cells, with other cell types expressing lower levels of the *Elmo1* transcript (**Fig. 1C**). During the bilateral ischemia/reperfusion induced acute kidney injury (IRI-AKI), *Elmo1* expression is transiently reduced in several cell types during the first 24 hours after injury, with recovery by 2 days following the initial insult (**Fig. 1C**). These data suggest that ELMO1 is linked to kidney disease and is expressed by kidney-resident cells as well as immune cells involved in acute kidney injury.

**Figure 1.**
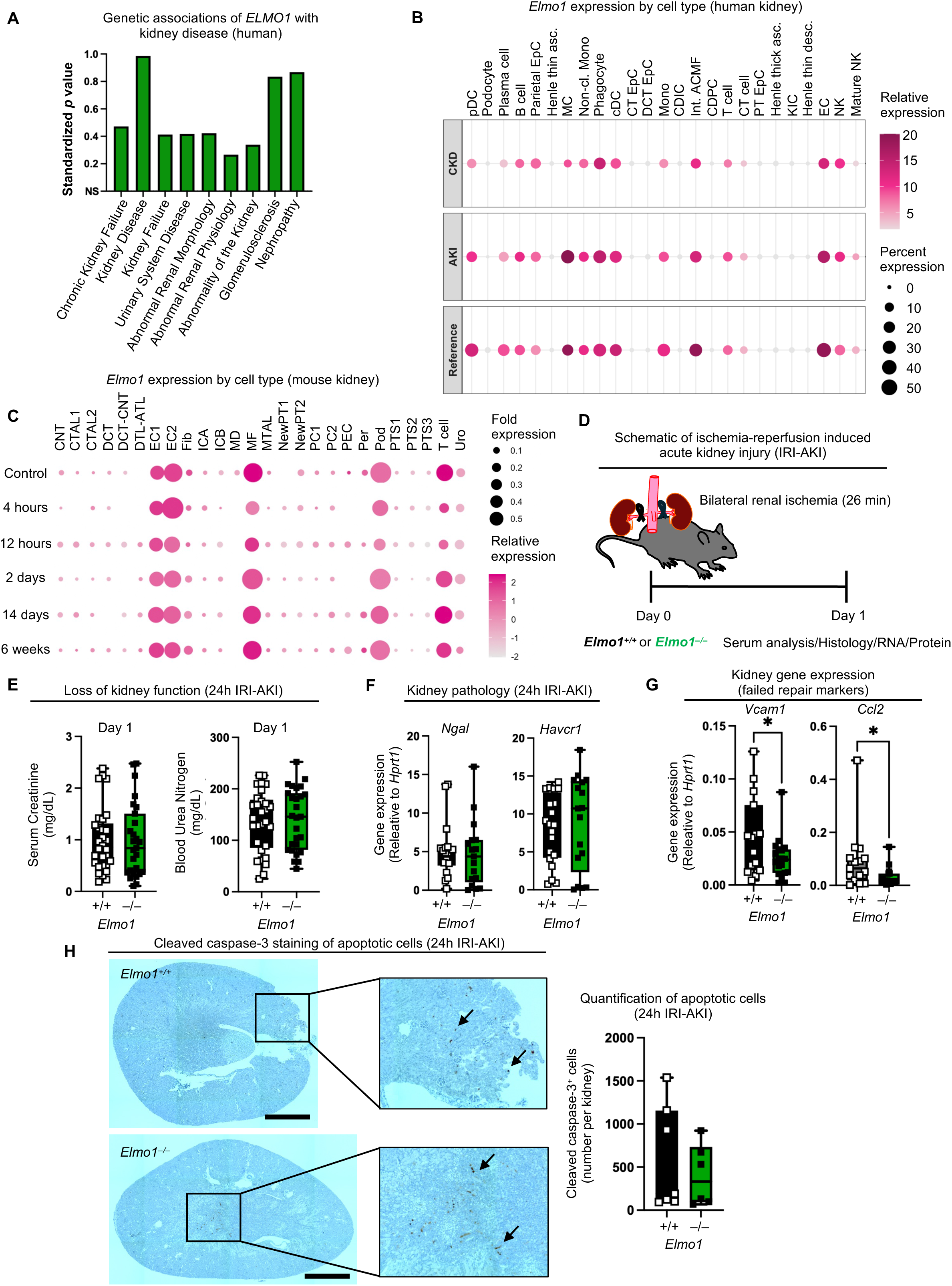
Ischemia-reperfusion induced kidney injury is not alleviated in *Elmo1^−/–^*mice. (**A**) Standardized *p* value plot of genetic associations of *ELMO1* with kidney disease in humans. (**B**) Single-cell RNAseq data of human kidneys in healthy (Reference) individuals and patients with acute kidney injury (AKI) or chronic kidney disease (CKD) were reanalyzed for *ELMO1* expression. Percent expression indicates the percentage of cells in each category with detectable *ELMO1,* and the color intensity indicates the relative expression level of *ELMO1* in the cell population, with darker color indicating higher expression level. (**C**) Single-nuclei RNAseq data of mouse ischemia-reperfusion injury (IRI) were reanalyzed for *Elmo1* transcript expression. Fold expression indicates the fraction of cells expressing *Elmo1*, and color intensity indicates relative expression level, as above. (**D**) Schematic representation of the bilateral kidney IRI model. (**E**) Serum creatinine and blood urea nitrogen (BUN) levels in *Elmo1*⁺/⁺ (n=29-30) and *Elmo1*⁻/⁻ (n=27) mice on Day 1 after IRI-AKI. Data are shown as mean ± SD; *p* values were determined by unpaired Student’s *t*-test; ns, not significant. Data shown are pooled from more than 3 independent experiments. Each symbol indicated an individual animal. (**F**) Kidney pathology following IRI-AKI, with relative gene expression of *Ngal* and *Havcr1* normalized to *Hprt1* in *Elmo1*^+/+^ (n=18-20) and *Elmo1*^−/–^ (n=17) mice. Data shown are pooled from more than 3 independent experiments. Each symbol indicated an individual animal. (**G**) Kidney gene expression of *Vcam1* and *Ccl2* relative to *Hprt1* in *Elmo1*^+/+^ (n=15) and *Elmo1*^−/–^ (n=19) mice on Day 1 after IRI-AKI. Data shown are pooled from more than 3 independent experiments. Each symbol indicated an individual animal. (**H**) Cleaved caspase-3 staining of apoptotic cells in kidney sections from *Elmo1*^+/+^ (n=7) and *Elmo1*^−/–^ (n=7) mice on Day 1 after IRI-AKI. Representative images are shown alongside quantification of positive cells per field. Scale bars indicate 1 mm. Data shown are pooled from two experiments. Data in panels D–F are shown as mean ± SD; p values were determined by unpaired Student’s *t*-test; \**p* < 0.05. pDC, plasmacytoid dendritic cell; EpC, epithelial cell; Henle thin asc., kidney loop of Henle thin ascending limb epithelial cell; MC, mast cell; Non-cl. Mono, non-classical monocyte; Phagocyte, mononuclear phagocyte; cDC, conventional dendritic cell; CT, kidney connecting tubule; DCT, kidney distal connecting tubule; Mono, monocyte; CDIC, kidney collecting duct intercalated cell; Int. ACMF, interstitial alternatively activated macrophage; CDPC, kidney collecting duct principal cell; CT cell, cytotoxic T cell; PT, proximal tubule; Henle thick asc., kidney loop of Henle thick ascending limb epithelial cell; KIC, kidney interstitial cell; Henle thin desc., kidney loop of Henle thin descending limb epithelial cell; EC, endothelial cell; NK, natural killer cell; CNT, connecting tubule; CTAL, thick ascending limb of loop of Henle in cortex; DCT, distal convoluted tubule; DTL, descending limb of loop of Henle; ATL, thin ascending limb of loop of Henle; EC, endothelial cells; Fib, fibroblasts; ICA, type A intercalated cells of collecting duct; ICB, type B intercalated cells of collecting duct; MD, macula densa; MF, macrophage; MTAL, thick ascending limb of loop of Henle in medulla; PT-S, S segment of proximal tubule; CPC, principle cells of collecting duct in cortex; PEC, parietal epithelial cells; Per, pericytes; Pod, podocytes; Uro, urothelium.

### Loss of ELMO1 does not alleviate tissue pathology in kidney ischemia-reperfusion injury (IRI-AKI)

Mice with the global deletion of *Elmo1* (*Elmo1*^−/–^) are generally healthy and fertile(35). At baseline, *Elmo1*^−/–^ mice have normal body and kidney weight and do not display elevation in the markers of kidney injury serum creatinine and blood urea nitrogen (BUN) or abnormalities in the kidney tissue architecture (**Supplementary Fig. 1**). Since changes in *Elmo1* expression levels were notable during the early stages of reperfusion (**Fig. 1C**), we next tested the role of ELMO1 in ischemia reperfusion induced acute kidney injury (IRI-AKI). We subjected *Elmo1*^−/–^ and control *Elmo1*^+/+^ mice to 26 minutes of bilateral ischemic renal injury, followed by reperfusion (**Fig. 1D**). Surprisingly, analysis of the serum-derived kidney disfunction markers after 24 hours of reperfusion revealed no significant differences between *Elmo1*^−/–^ and control mice (**Fig. 1E**). Similarly, no differences were observed in transcript levels of kidney injury markers *Ngal* and *Havcr1* (which encodes Kim1, **Fig. 1F**), or the overall tissue pathology based on hematoxylin and eosin staining (**Supplementary Fig. 2A**), suggesting that the initial renal injury in IRI-AKI is not impacted by the absence of ELMO1. We also quantified early markers of failed renal repair, *Vcam1* and *Ccl2*, associated with pathological remodeling during renal tubular repair(42). Increase in *Vcam1* and *Ccl2* transcripts was significantly lower in *Elmo1*^−/–^ IRI-AKI kidneys, compared to control (**Fig. 1G**), suggesting that ELMO1 could decrease renal repair.

ELMO1 can regulate the inflammatory response that contributes to maladaptive repair after renal injury(42,43), so we next analyzed inflammatory cytokines in the IRI-AKI kidneys. Gene expression analysis demonstrated trends toward reduced expression of transcripts that encode several inflammatory cytokines in *Elmo1*^−/–^ IRI-AKI kidneys, although these differences did not reach statistical significance (**Supplementary Fig. 2B**). IL-6 and TNF-α contribute to post-ischemic sequalae(44,45), and we found that IL-6 is significantly reduced in *Elmo1*^−/–^ IRI-AKI kidneys (local inflammation), compared to *Elmo1*^+/+^ mice (**Supplementary Fig. 2C**). Importantly, IL-6 and TNF-α were completely absent from the serum of *Elmo1*^−/–^ IRI-AKI mice (systemic inflammation), whereas high levels of both cytokines could be detected in several *Elmo1*^+/+^ IRI-AKI controls (**Supplementary Fig. 2D**). Overall, these data suggest that ELMO1 contributes to the systemic inflammatory response that follows the ischemic insult and might promote maladaptive repair of the injured kidney after reperfusion.

Recruitment of neutrophils during ischemia-reperfusion critically contributes to the pathogenesis of IRI-AKI(46). Since ELMO1 promotes neutrophil migration into tissues(34,47), we next tested if neutrophil infiltration or activation in IRI requires ELMO1. We first asked if neutrophils contribute to the systemic inflammation in this model by quantifying serum levels of neutrophil granule-stored enzymes, neutrophil elastase, and myeloperoxidase, by ELISA. As noted for inflammatory cytokines (**Supplementary Fig. 2D**), several *Elmo1*^+/+^ IRI-AKI mice had high systemic levels of the neutrophil enzymes, which were not observed in any of the *Elmo1*^−/–^ IRI-AKI mice (**Supplementary Fig. 2E**). However, no significant differences between *Elmo1*^+/+^ and *Elmo1*^−/–^ IRI-AKI mice were observed in the levels of neutrophil elastase and myeloperoxidase in the kidney tissue (**Supplementary Fig. 2F**), and quantification of kidney areas positive for Ly6G, a neutrophil-specific marker, revealed no differences in the neutrophil influx into the kidneys (**Supplementary Fig. 2G**). These data suggest that the neutrophil influx into the IRI-AKI kidneys does not require ELMO1, but that neutrophil activation and release of the granule contents into systemic circulation may be hastened by *Elmo1* expression.

ELMO1 was originally identified as a promoter of apoptotic cell clearance by phagocytes(33). Since renal cell death directly affects AKI outcomes(21,23), we stained kidney sections from control and *Elmo1*^−/–^ mice subjected to IRI-AKI with an antibody specific for a marker of apoptosis, cleaved caspase-3. We noted a variable and generally low number of cleaved caspase-3-stained apoptotic cells, which was comparable between *Elmo1*^+/+^ and *Elmo1*^−/–^ mice (**Fig. 1H**). This is consistent with the notion that this model of early ischemia-reperfusion kidney injury associates with non-apoptotic programmed cell death(23). Collectively, these data suggest that ELMO1 does not contribute to tissue pathology in early ischemic kidney injury.

### Loss of ELMO1 aggravates tissue injury and impairs removal of apoptotic cells in cisplatin-induced acute kidney injury (cisplatin-AKI)

Apoptosis is the predominant type of cell death observed during kidney injury induced by nephrotoxic agents such as cisplatin(14). To address if loss of ELMO1 influences disease pathology in a model of kidney injury in which extensive apoptosis occurs, we subjected *Elmo1*^−/–^ mice to nephrotoxic injury caused by cisplatin injection (cisplatin-AKI, **Fig. 2A**). We first quantified the progressive loss of kidney function by analyzing serum creatinine and BUN on days 2 and 3 after cisplatin injection. Although serum creatinine levels did not differ between *Elmo1*^+/+^ and *Elmo1*^−/–^ mice in cisplatin-AKI, we noted significantly elevated BUN levels in *Elmo1*^−/–^ mice compared to controls on day 2 post cisplatin injection, with a trend in increase that persisted on day 3 (**Fig. 2B**). Gene expression analysis of the kidney injury markers *Ngal* and *Havcr1* did not reveal significant differences (**Fig. 2C**), suggesting that the extent of kidney injury is not affected by the loss of ELMO. In addition, inflammatory cytokine gene expression was low in the kidneys of all mice (**Supplementary Fig. 3**). However, kidney tissue damage was significantly accelerated in *Elmo1*^−/–^ mice, compared to controls (**Fig. 2D**).

**Figure 2.**
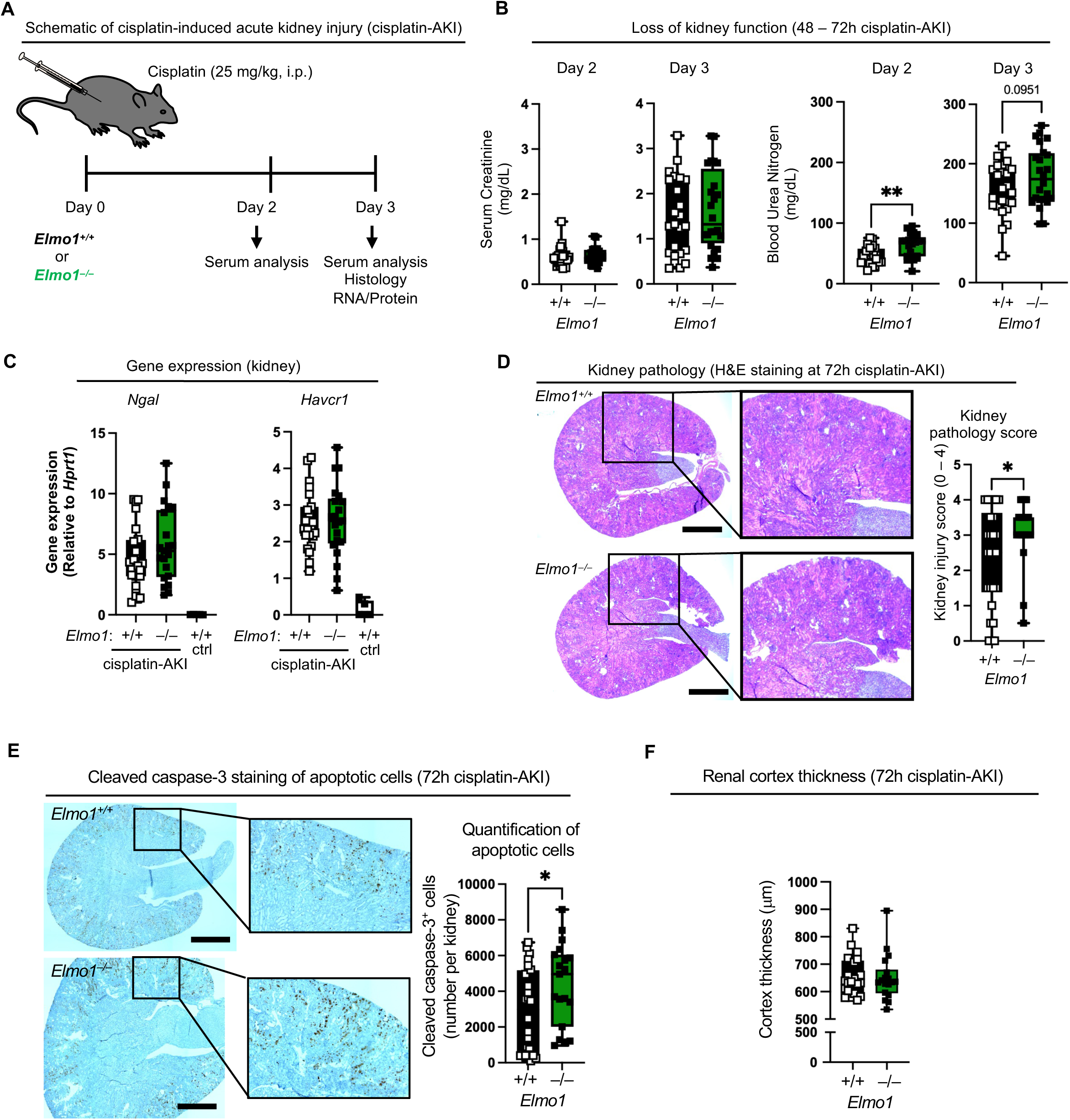
Loss of ELMO1 aggravates cisplatin-induced acute kidney injury (cisplatin-AKI) in mice. (**A**) Schematic representation of the cisplatin-induced acute kidney injury (AKI) model. (**B**) Serum creatinine and blood urea nitrogen (BUN) levels in the serum of *Elmo1*^+/+^ (n=24) and *Elmo1*^−/–^ (n=24) mice on day 2 and day 3 after cisplatin injection. Data shown are pooled from 3 independent experiments. (**C**) Kidney pathology following cisplatin-induced AKI, with relative gene expression of *Ngal* and *Havcr1* normalized to *Hprt1* in *Elmo1*^+/+^ (n=24), *Elmo1*^−/–^ (n=21), and control (ctrl, n=5) mice at day 3 post-injection. Data are pooled from 3 independent experiments. Each symbol indicated an individual animal. (**D**) Kidney pathology at 72 h post cisplatin-induced AKI, assessed by hematoxylin-eosin (H&E) staining. Representative images are shown alongside quantification of kidney pathology scores (0–4) in *Elmo1*^+/+^ (n=26) and *Elmo1*^−/–^ (n=21) mice. Scale bars indicate 1 mm. Data are pooled from 3 independent experiments. Each symbol indicated an individual animal. (**E**) Representative images of cleaved caspase-3 staining of apoptotic cells in kidney sections from *Elmo1*^+/+^ (n=26) and *Elmo1*^−/–^ (n=19) mice at 72 h post cisplatin-induced AKI. Scale bars indicate 1 mm. Data are pooled from 3 independent experiments. Each symbol indicated an individual animal. (**F**) Renal cortex thickness in μm in mice at 72 h post cisplatin-induced AKI. Each symbol indicates an individual animal. Data are shown as mean ± SD; p values were determined by unpaired Student’s *t*-test; \**p* < 0.05, \*\**p* < 0.01.

Since renal cell death directly affects AKI outcomes(21,23), and ELMO1 can mediate the removal of apoptotic cells in tissues(35,48), we next quantified cleaved caspase-3-stained apoptotic cells in cisplatin-AKI. Apoptotic cells were abundant in the kidneys of mice subjected to cisplatin-AKI, with loss of ELMO1 resulting in significantly increased total number of apoptotic cells (**Fig. 2E**), correlating with accelerated kidney pathology and elevated BUN in the serum (**Fig. 2D**). However, apoptotic cell accumulation in the kidneys of *Elmo1*^−/–^ mice did not result in a significant increase in the renal cortex thickness (**Fig. 2F**), possibly suggestive of compensatory, ELMO1-independent, efferocytosis pathways.

Collectively, these data suggest that increased nephrotoxic injury in ELMO1 deficient mice could be mediated through a mechanism that does not involve regulation of the inflammatory response but likely involves cell death and/or clearance modulation.

### ELMO1 is not required for renal tubular epithelial cell (RTEC) death or efferocytosis

To address if increase in cleaved caspase-3-stained apoptotic cells in the kidneys of *Elmo1*^−/–^ mice during cisplatin-AKI (**Fig. 2E**) could be due to increased susceptibility of renal cells to cisplatin-induced death, we established primary cultures of renal tubular epithelial cells (RTEC). *Elmo1*^+/+^ and *Elmo1*^−/–^ RTEC cultures did not contain CD45^+^ cells, were 80-90% enriched for the epithelial cell adhesion molecule EpCAM (**Supplementary Fig. 4A-B**) and expressed the tight junction marker ZO-1 (**Supplementary Fig. 4C**). We verified ELMO1 expression in primary RTECs by immunoblot analysis (**Supplementary Fig. 4D**). Next, primary RTEC cultures were treated with increasing concentrations of cisplatin for 2 days, and apoptosis was assessed by flow cytometry. We noted cisplatin-induced increase in the percentage of apoptotic RTECs (AnnexinV^+^7AAD^−^) at cisplatin doses above 50 μM; however, no differences in the amount of apoptosis were observed between *Elmo1*^+/+^ and *Elmo1*^−/–^ RTEC cultures (**Supplementary Fig. 4E-F**). These data suggest that increased apoptotic cells in the kidneys of cisplatin-treated *Elmo1*^−/–^mice are likely not due to increased susceptibility to cell death.

Next, we addressed whether loss of ELMO1 impairs phagocytosis of apoptotic cells present during the kidney pathology. In the kidney, tubular epithelial cells have been shown to function as phagocytes after acute renal injury(49), so we tested whether *Elmo1*^−/–^ RTEC-mediated efferocytosis is impaired. As efferocytosis targets, we first used Jurkat cells that have been rendered apoptotic by ultraviolet (UV) light exposure (**Fig. 3A**, see Methods). Apoptotic Jurkat cells were stained with the pH-sensitive dye CypHer5E, which becomes fluorescent when exposed to the acidic environment of the phagolysosome, and were fed to primary RTECs established from *Elmo1*^+/+^ and *Elmo1*^−/–^ mice (**Fig. 3B**). Although the percentage of CypHer5E^+^ RTECs that ingested one of more apoptotic Jurkat cells did not differ between *Elmo1*^+/+^ and *Elmo1*^−/–^ RTECs, we noted a strong trend toward the reduced number of engulfed cells per *Elmo1*^−/–^ RTEC (compared to control), as evident by the reduced geometric mean fluorescence index (MFI) of the CypHer5E signal (**Fig. 3C**). We also tested RTEC efferocytosis of cisplatin-killed RTECs labeled with CypHer5E. Again, the percentage of CypHer5E^+^ RTEC did not depend on ELMO1, and reduced CypHer5E MFI in *Elmo1*^−/–^ RTECs did not reach significance (**Fig. 3D**). Overall, these data suggest that loss of ELMO1 does not significantly impact renal epithelial cell death or efferocytosis.

**Figure 3.**
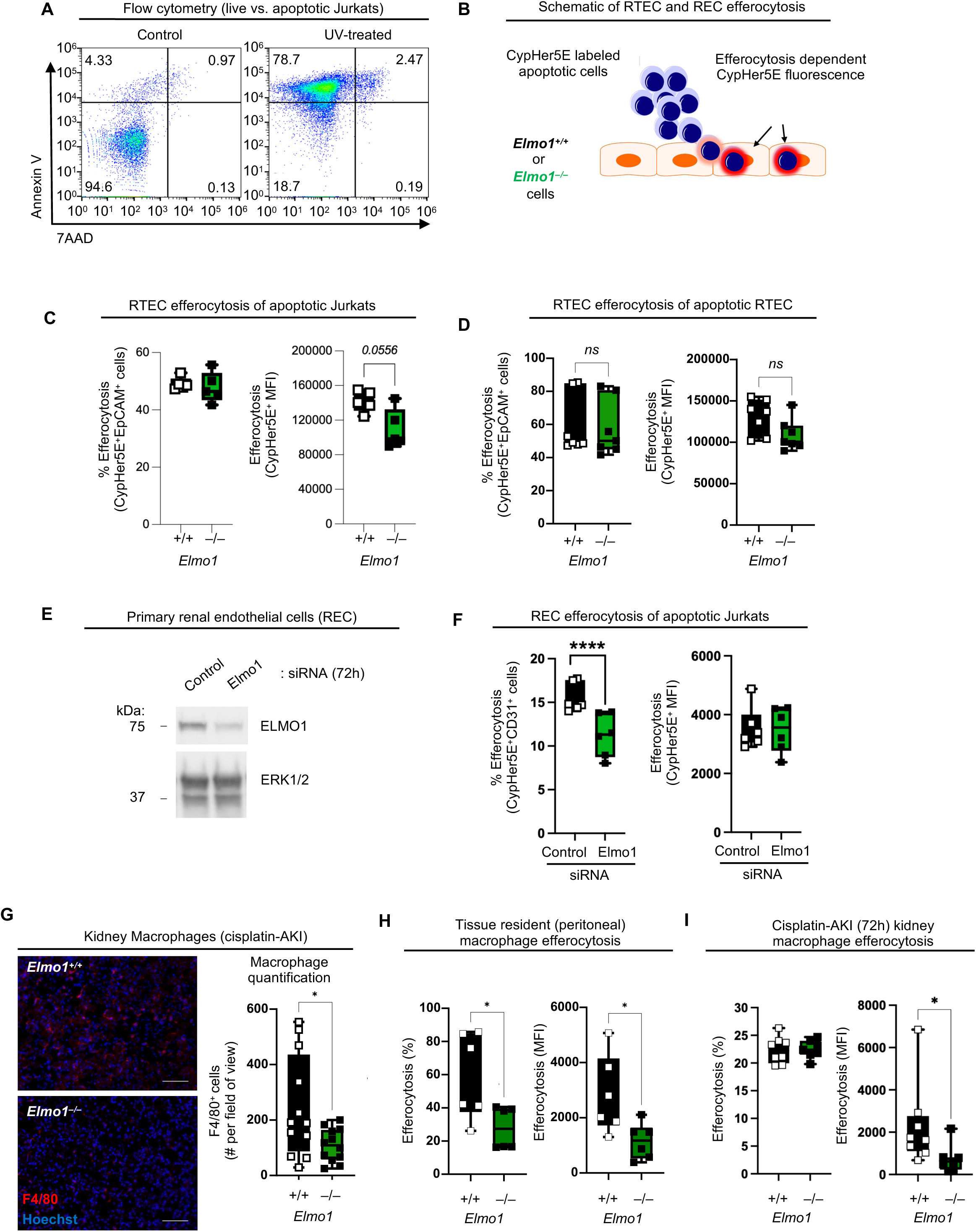
ELMO1 contribution to efferocytosis in renal epithelial and endothelial cells and macrophages. (**A**) Flow cytometry analysis of Annexin V and 7AAD staining in Jurkats cells. Representative plots of control/live and 2 hours after UV exposure cells are shown, from more than 3 independent experiments. (**B**) Schematic representation of efferocytosis assays. CypHer5E-stained apoptotic target cells become fluorescent after engulfment by phagocytes due to the sensitivity of the CypHer5E dye to acidic pH present in phagolysosomes. (**C**) Efferocytosis of UV-induced apoptotic Jurkat cells by RTEC derived from *Elmo1*^+/+^ (n=5) and *Elmo1*^−/–^ (n=5) mice. Graphs show percentage of CypHer^+^EpCAM^+^ RTEC and mean fluorescence intensity (MFI) of CypHer5E in the CypHer^+^EpCAM^+^ RTEC after 2 hours of efferocytosis. Data are pooled from 2 independent experiments. (**D**) Efferocytosis of cisplatin-induced apoptotic RTEC by RTEC from *Elmo1*^+/+^ (n=7) and *Elmo1*^−/–^ (n=7) mice. Graphs show percentage of CypHer^+^EpCAM^+^ RTEC and mean fluorescence intensity (MFI) of CypHer5E in the CypHer^+^EpCAM^+^ RTEC after 2 hours of efferocytosis. Data are pooled from 2 independent experiments. (**E**) Primary renal endothelial cells (REC) 72 hours after transfection with Elmo1-targeting or Control siRNA. Immunoblot analysis shows reduced ELMO1 protein levels in Elmo1 siRNA transfected cells. ERK1/2 was used for protein loading control. (**F**) Efferocytosis of UV-induced apoptotic Jurkat cells by REC. Graphs show percentage of CypHer^+^CD31^+^ REC and mean fluorescence intensity (MFI) of CypHer5E in the CypHer^+^CD31^+^ REC after 2 hours of efferocytosis. Data are pooled from 2 independent experiments, with n=6. (**G**) Kidney sections from *Elmo1*^+/+^ (n=12) and *Elmo1*^−/–^ (n=12) mice 72 hours after cisplatin-AKI were stained for macrophages (F4/80). Hoechst was used to stain the cell nuclei. Scale bars indicate 100 µm. (**H**) Efferocytosis of UV-induced apoptotic thymocytes by tissue-resident peritoneal macrophages from *Elmo1*^+/+^ (n=6) and *Elmo1*^−/–^ (n=6) mice. Graphs show percentage of CypHer^+^CD11b^+^F4/80^+^ macrophages and mean fluorescence intensity (MFI) of CypHer5E in the CypHer^+^CD11b^+^F4/80^+^ macrophages after 1 hour of efferocytosis. Data are pooled from two experiments. (**I**) Efferocytosis of UV-induced apoptotic thymocytes by renal macrophages isolated after 72 h of cisplatin-AKI from *Elmo1*^+/+^ (n=8) and *Elmo1*^−/–^ (n=7) mice. Graphs show percentage of CypHer^+^F4/80^+^ macrophages and mean fluorescence intensity (MFI) of CypHer5E in the CypHer^+^F4/80^+^ macrophages after 1 hour of efferocytosis. Data are from one experiment, with each symbol indicating an individual animal. Data in panels C-I are shown as mean ± SEM; p values were determined by unpaired Student’s *t*-test with Mann-Whitney correction; \**p* < 0.05. **** *p* < 0.0001. ns, not significant.

### ELMO1 promotes clearance of apoptotic cells by endothelial cells and macrophages

ELMO1 is robustly expressed in the kidney endothelial cells, macrophages, podocytes, and T cells (**Fig. 1B-C**). Since T cell phagocytosis has not been reported, and since cleaved caspase-3-stained apoptotic cells did not appear to overlap with podocyte-enriched glomeruli in either *Elmo1*^+/+^ or *Elmo1*^−/–^ cisplatin-AKI kidneys (**Supplementary Fig. 5**, arrows point to areas of the kidney consistent with glomerular structures), we focused on endothelial cells and macrophages as phagocytes in which ELMO1 could promote efferocytosis.

We obtained primary renal endothelial cells (REC) purified from C57BL/6 mice and knocked down ELMO1 using an siRNA approach (see Methods). We verified reduced ELMO1 protein levels in Elmo1 siRNA-treated REC, compared to control siRNA, by immunoblot analysis (**Fig. 3E**). Next, we tested if reduced ELMO1 impacts REC efferocytosis of apoptotic Jurkat cells. We noted a significantly reduced percentage of CypHer5E^+^ REC after siRNA mediated knockdown of Elmo1, compared to control (**Fig. 3F**), suggesting that ELMO1 could contribute to efferocytosis in renal endothelial cells.

Macrophages are professional phagocytes that are highly capable of performing efferocytosis. We first tested whether the macrophage number in the kidneys of cisplatin-treated mice is influenced by the loss of ELMO1. *Elmo1*^+/+^ and *Elmo1*^−/–^ cisplatin-AKI kidneys were stained with the macrophage marker F4/80, and macrophage numbers quantified by immunofluorescence. We observed a significantly reduced number of macrophages in the kidneys of *Elmo1*^−/–^ cisplatin-AKI mice, compared to *Elmo1*^+/+^ (**Fig. 3G**). We next tested if ELMO1 deficiency can reduce macrophage efferocytosis using peritoneal macrophages as a model for tissue resident macrophages, and CypHer5E-labeled apoptotic thymocytes as a model target. *Elmo1*^−/–^ macrophages had significantly reduced efferocytosis compared to controls, with both the percentage of macrophages that engulfed one or more targets and the number of targets per macrophage reduced (**Fig. 3H**). Finally, we also tested efferocytosis of renal macrophages purified from the kidneys of cisplatin-injected *Elmo1*^+/+^ and *Elmo1*^−/–^ mice *ex vivo*. Although the percentage of efferocytotic macrophages did not differ between *Elmo1*^+/+^ and *Elmo1*^−/–^ renal macrophages, *Elmo1*^−/–^ macrophages engulfed significantly fewer targets per cell (**Fig. 3I**). Together, these data suggest that ELMO1 promotes renal macrophage recruitment and/or expansion in cisplatin-AKI and could mediate macrophage efferocytosis.

### Deletion of ELMO1 in macrophages does not recapitulate global loss of ELMO1 in cisplatin-AKI

Because we noted reduced macrophage numbers in cisplatin treated *Elmo1*^−/–^ mice and reduced macrophage mediated efferocytosis, we next tested if macrophage ELMO1 protects the kidneys from cisplatin-AKI. To do this, we crossed *Elmo1^fl/fl^* mice(35) with a macrophage-specific Cre mouse line (*Csf1r*-Cre(50)). We verified the loss of ELMO1 protein from *Elmo1^fl/fl^Csf1r*-Cre macrophages, compared to *Elmo1^fl/fl^* controls by immunoblot analysis (**Fig. 4A**). Next, we subjected *Elmo1^fl/fl^* and *Elmo1^fl/fl^Csf1r*-Cre mice to cisplatin-AKI (**Fig. 4B**). Compared to *Elmo1^fl/fl^* mice, *Elmo1^fl/fl^Csf1r*-Cre mice did not exhibit significantly elevated serum creatinine or BUN (**Fig. 4C**) and did not have significant increases in expression of kidney injury markers *Ngal* and *Havcr1* (**Fig. 4D**) or renal tissue pathology (**Fig. 4E**). Finally, cisplatin injected *Elmo1^fl/fl^Csf1r*-Cre mice did not have reduced numbers of renal macrophages (**Fig. 4F**) or significantly increased cleaved caspase-3 stained apoptotic cell numbers in the kidneys (**Fig. 4G-H**).

**Figure 4.**
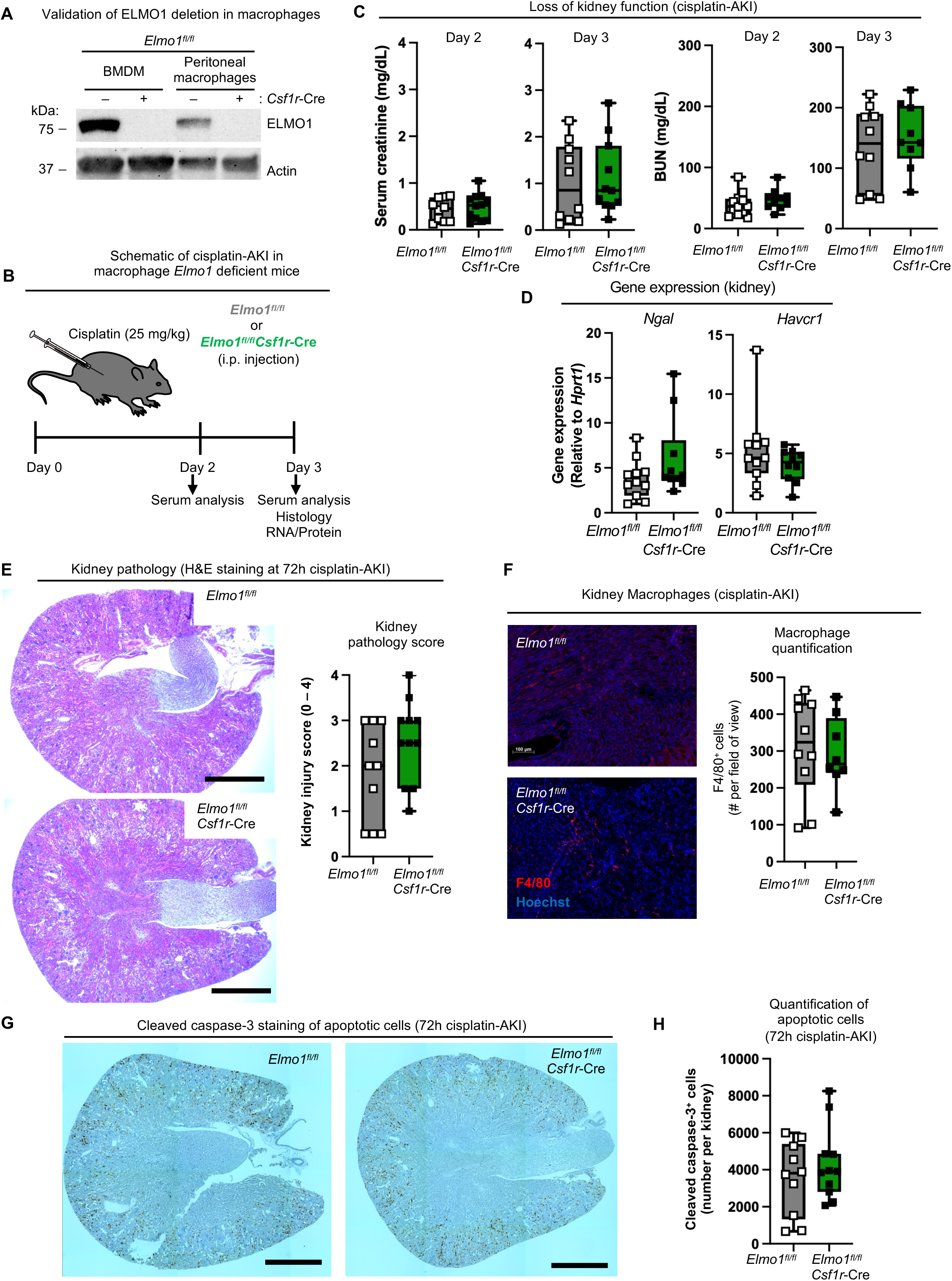
Macrophage-specific *Elmo1* deletion is not sufficient to recapitulate the global loss of ELMO1 in cisplatin-AKI. (**A**) Immunoblot analysis of ELMO1 protein levels in bone marrow-derived macrophages (BMDM) and peritoneal macrophages from *Elmo1^fl/fl^* and *Elmo1^fl/fl^Csf1r-Cre* mice. Actin was used as protein loading control. Representative of two independent experiments. (**B**) Schematic representation of cisplatin-AKI in mice with macrophage-specific deletion of *Elmo1*. (**C**) Serum creatinine and blood urea nitrogen (BUN) serum levels in *Elmo1^fl/fl^*(n=10-12) and *Elmo1^fl/fl^Csf1r-Cre* (n=9-12) mice on day 2 and day 3 post-injection. (**D**) Kidney pathology following cisplatin-induced AKI, with relative gene expression of *Ngal* and *Havcr1* normalized to *Hprt1* in *Elmo1^fl/fl^*(n=11) and *Elmo1^fl/fl^Csf1r-Cre* (n=10) mice on day 3 post-injection. (**E**) Kidney pathology in *Elmo1^fl/fl^* (n=10) and *Elmo1^fl/fl^Csf1r-Cre* (n=11) mice on day 3 post cisplatin-induced AKI, assessed by hematoxylin-eosin (HE) staining. Representative images are shown. Scale bars indicate 1 mm. Quantification of kidney pathology scores (0–4) is shown on the right. (**F**) Kidney sections from *Elmo1^fl/fl^* (n=10) and *Elmo1^fl/fl^Csf1r-Cre* (n=8) mice 72 hours after cisplatin-AKI were stained for macrophages (F4/80). Hoechst was used to stain the cell nuclei. Scale bar indicates 100 µm. (**G**) Cleaved caspase-3 staining of apoptotic cells in kidney sections from *Elmo1^fl/fl^* (n=10) and *Elmo1^fl/fl^Csf1r-Cre* (n=11) mice on day 3 post cisplatin-induced AKI. Representative images are shown. Scale bars indicate 1 mm. (**H**) Quantification of apoptotic cells per field from panel G. Data in panels C–H are shown as mean ± SD; p values were determined by unpaired Student’s *t*-test. Data shown are pooled from 2 independent experiments. Each symbol indicated an individual animal.

Collectively, these data show that macrophage-specific deletion of *Elmo1* in *Elmo1^fl/fl^Csf1r*-Cre mice does not recapitulate the global ELMO1 deficiency in cisplatin-AKI, suggesting that ELMO1 likely contributes to efferocytosis in multiple renal cell types.

## Discussion

Our findings presented here reveal a context-dependent role of ELMO1 in acute kidney injury, integrating insights from human genetics and mouse models. We found that genetic associations between *ELMO1* and kidney pathology in humans extend beyond diabetic nephropathy and that diverse renal cell types express ELMO1 in human and mouse kidneys.

In the mouse model of kidney ischemia-reperfusion injury (IRI-AKI), we surprisingly found that global *Elmo1* deletion did not alter early tissue injury, as assessed by serum creatinine, BUN, histology, or cleaved caspase-3 staining. However, in *Elmo1^−/–^* mice, we noted trends toward reduced systemic and local inflammation at early time points after injury, suggesting that loss of ELMO1 could affect neutrophil activation and degranulation rather than recruitment, since neutrophil influx was not reduced in the kidneys of *Elmo1^−/–^* mice following IRI. This contrasts with inflammatory arthritis models, where ELMO1 is critical for neutrophil tissue infiltration(34). Such divergence could suggest tissue-specific ELMO1 requirement in neutrophil recruitment and highlights that systemic cytokine release can be decoupled from local cell influx. We also noted decreased early renal expression of *Vcam1* and *Ccl2*, possibly suggestive of reduced inflammatory tissue activation associated with maladaptive renal remodeling(43). Whether recovery of *Elmo1*^−/–^ mice at the later time points after ischemic injury could be enhanced remains to be addressed.

By contrast, in cisplatin-induced nephrotoxic AKI, *Elmo1* deficiency significantly worsened early injury, with elevated BUN, accelerated histopathology, and increased numbers of uncleared apoptotic cells. We found that early cisplatin-AKI is dominated by apoptosis, whereas the inflammatory response in cisplatin-AKI was mild and not dependent on ELMO1, suggesting that ELMO1 function in cisplatin-AKI likely relates to its role in apoptotic cell clearance, or efferocytosis. Engineered efforts to enhance efferocytosis through expression of a chimeric receptor that constitutively couples the activity of the efferocytosis receptor BAI-1 with ELMO1-mediated cell signaling (BAI-1/ELMO1 fusion receptor, or BELMO) have shown promise in renal injury protection(51). However, ELMO1 function is not unique to BAI-1 signaling(52–54) and likely plays a broader role in efferocytosis regulation.

In the mouse kidney, we noted that many cell types express the *Elmo1* transcript, with macrophages, endothelial cells, and podocytes particularly enriched. Although *Elmo1* expression was generally low in the subsets of renal epithelial cells, these cells are the primary renal target of cisplatin(14). In addition, healthy renal tubular epithelial cells perform efferocytosis to clear damaged tubules(49), and BELMO expression in renal epithelial cells (via *Pepck*-Cre) was required for efferocytosis-mediated renal protection in AKI(51). We first addressed whether the loss of ELMO1 might increase susceptibility of primary renal tubular epithelial cells (RTEC) to cisplatin-induced death in culture; however, no differences in cell death were observed between ELMO1 sufficient and deficient cells exposed to a broad range of cisplatin concentrations. We also addressed if the efferocytosis function of RTECs requires ELMO1. Although we noted a trend toward the reduced per cell *Elmo1^−/–^* RTEC uptake of apoptotic corpses, this difference was small and did not reach statistical significance, suggesting that loss of ELMO1 efferocytosis function in renal epithelial cells is not likely to be responsible for the increased renal apoptotic cell burden in cisplatin-AKI.

Because we did not observe an overlap between the cleaved caspase-3-stained apoptotic cells and the podocyte-residing renal glomeruli, we addressed the role of ELMO1 in endothelial cell- and macrophage-mediated efferocytosis. siRNA-based knockdown of Elmo1 in purified primary renal endothelial cells (REC) resulted in a significantly reduced percentage of efferocytosis, without an impact on the uptake per cell, possibly reflecting the incomplete loss of ELMO1 on a population basis. The efficiency of REC efferocytosis was low, with only 10-15% of the cells engulfing one or more corpses, again suggesting a reduced likelihood that REC are the primary renal phagocytes in cisplatin-AKI. On the other hand, macrophages are professional phagocytes with a high efferocytosis capacity(55). We observed reduced macrophage numbers in the cisplatin-AKI kidneys of *Elmo1^−/–^* mice and significantly reduced efferocytosis in the ELMO1-deficient tissue resident macrophages and cisplatin-AKI renal macrophages. Therefore, we addressed the role of macrophage-specific *Elmo1* deletion in cisplatin-AKI. Surprisingly, cisplatin treatment of mice with *Csf1r*-Cre mediated *Elmo1* deletion, despite effectively ablating ELMO1 protein in both tissue resident and monocyte-derived macrophages, did not recapitulate the global loss of ELMO1. Although we noted trends in elevated BUN, tissue pathology and apoptotic cell burden of *Elmo1^fl/fl^Csf1r*-Cre cisplatin-AKI mice, none reached significance, suggesting that ELMO1 could contribute to macrophage function in cisplatin-AKI, but that its role in renal efferocytosis likely requires contribution from additional cell types.

Through formation of a complex with DOCK family proteins, ELMO1 functions as a guanine exchange factor (GEF) for Rac GTPases to promote actin cytoskeleton rearrangements in phagocytosis and cell migration (33). Rac1 has been shown to play a context-dependent and cell type-specific role in various models of renal injury(56–60). Whether loss of ELMO1 GEF activity would phenocopy the global deficiency of the ELMO1 protein in cisplatin-AKI remains to be determined.

Genome-wide association studies link *ELMO1* gene with diabetic nephropathy in diverse human populations(36–39). In the Akita mouse model of diabetic nephropathy, elevated ELMO1 levels were shown to associate with enhanced renal disease, whereas reduced ELMO1 offered protection(61), consistent with our notion that ELMO1 has a complex role in renal pathology. ELMO1 could contribute to some facets of the early inflammatory response in ischemic injury, whereas in cisplatin-mediated nephrotoxic injury, ELMO1 could promote efferocytosis to reduce the renal dying cell burden. Although we did not note accumulation of apoptotic cells in early ischemia-reperfusion, it remains to be addressed if apoptotic cells can be observed at later stages and if ELMO1 could contribute to their removal in this model. Similarly, our data also highlight the difference between endogenous ELMO1 and the engineered BELMO (in which ELMO1 is fused to the BAI-1 receptor). Whether ELMO1 over-expression in renal epithelial cells would offer protection during renal injury is not known. Importantly, macrophage-specific BELMO-induced efferocytosis boost was not sufficient for renal protection(51), and reduced efferocytosis in primary *Elmo1^−/–^* macrophages has not been reported in previous studies(35,62,63), likely due to differences in culture conditions, suggesting possible compensation strategies (such as ELMO2 upregulation(35)) that macrophages employ to maintain efficient efferocytosis function. Collectively, our data presented here uncover a complex and context-dependent role for ELMO1 in acute renal injury.

## Methods

### Mice

*Elmo1^fl/fl^* and *Elmo1^−/–^* mice have been described previously(35). To generate mice with deletion of *Elmo1* in the macrophage lineage, *Elmo1^fl/fl^* mice were crossed to *Csf1r*-Cre mice (Jackson mice #034470(50). Age- and sex-matched littermate control animals were used for all experiments. Animals are housed in Allentown ventilated racks with reverse osmosis water through an automatic watering system. They receive ad libitum diet and are kept in a 14:10 light: dark cycle at 72 degrees F and 40% humidity. All animal procedures were approved by and performed according to the guidelines of the Institutional Animal Care and Use Committee (IACUC) at the University of Virginia under the protocol #4320.

### Kidney Injury

Surgical procedures involving clamping both renal pedicles for 26 minutes were performed as previously described(64). After removing the clamps, the kidneys were allowed to reperfuse for 24 hours before euthanasia. For IRI surgical procedures, mice were anesthetized with an intraperitoneal injection of ketamine (120 mg/kg) and xylazine (12 mg/kg). Buprenorphine (0.15 mg/kg) was used as an analgesic, subcutaneously injected immediately after surgery. For cisplatin experiments, mice were intraperitoneally injected with cisplatin (25 mg/kg) and sacrificed 72 h after the cisplatin injection. The blood samples and kidney tissues were collected for further analysis at indicated time points. Serum separated from the collected blood was used for serum creatinine (SerCr) and blood urea nitrogen (BUN) analysis. Serum creatinine was measured using an enzymatic method kit (Diazyme Laboratories; Poway, CA, US) and BUN using a colorimetric assay kit (Arbor Assays; Ann Arbor, MI, US) according to the manufacturer’s instructions.

### Histology

Kidney tissue of mice was fixed in 10% neutral buffered formalin for 24 hours. Hematoxylin and eosin (H&E) staining was performed at the University of Virginia Research Histology Core. Histopathology scoring was performed by an investigator blinded to the mouse genotypes. For the kidney injury scoring, the following criteria were used: 0, no injury; 1, <25% damage; 2, 25-50% damage; 3, 50-75% damage; 4, >75% damage. Cleaved caspase-3 staining was performed at the University of Virginia Biorepository and Tissue Research Facility.

### Immunofluorescence

Kidney tissue of mice was fixed in periodate-lysine-paraformaldehyde (PLP) fixative for 24 h, washed with PBS, and equilibrated in 18% sucrose solution overnight. Fixed kidneys were next embedded and frozen in Optimal Cutting Temperature (OCT) compound (Sakura; Turrance, CA, US). Tissue was sectioned 5 μm thick, permeabilized with 0.5% Triton X-100 (Fisher; Waltham, MA, US), and blocked using 3% bovine serum albumin. Tissue sections were incubated with anti-Ly6G-PE (1:200, BioLegend #127608; San Diego, CA, US) in the dark for 1 h. For RTECs staining, cells were fixed with 4% PFA (EMS; Cleveland, OH, US), permeabilized with 0.1% Triton X-100 (Fisher), and blocked using 3% bovine serum albumin. Then, the cells were incubated with anti-ZO-1 antibody (1:500, eBioscience #14977637; San Diego, CA, US) overnight and with a secondary antibody (1:1000) for 1 h. All the samples were mounted using ProLong Gold antifade reagent (Invitrogen; Carlsbad, CA, US) containing DAPI counterstain. Images were acquired using either the Carl Zeiss Axiovert 200 microscopy system with Apotome imaging and Axiovision software (Zeiss; Oberkochen, Germany) or an EVOS FL Auto (Thermo Fisher; Waltham, MA, US) and integrated system.

### Gene Expression Analysis

For RNA isolation, the kidney tissue was placed in DNA/RNA Shield (Zymo Research; Irvine, CA, US). Total RNA from kidney tissue was isolated using Quick-RNA MiniPrep kit (Zymo Research) and complementary DNA was prepared using iScript cDNA synthesis kit (BioRad; Hercules, CA, US) according to the manufacturer’s instructions. Quantitative gene expression for target and housekeeping genes was done using Taqman probes (Thermo Fisher) run on a CFX Connect Real-Time System (BioRad).

### Isolation and culture of primary mouse RTECs

The kidneys were minced with collagenase type I (1.5 mg/ml; Sigma), diluted with RPMI (Genesee Scientific; El Cajon, CA, US). After incubation for 60 min at 37 ^°^C, the digested kidneys were pushed through a sterile 70-μm nylon cell strainer (Corning; Corning, NY, US) and collected in a 50 mL tube. Kidney cells were centrifuged at 2,000 rpm for 10 min, and the red blood cells were removed with Hybri-Max Red Blood Cell Lysing Buffer (Sigma, St Louis, MO, US). After centrifugation at 2,000 rpm for 10 min, the pellet was collected and resuspended in 10% FBS-RPMI with added EGF (10 ng/ml, Peprotech; Cranbury, NJ, US). The cells were plated onto 100-mm culture dishes and left unstirred in an incubator for 48 hours. EpCAM^+^ RTECs were obtained using immunomagnetic beads coated with PE-conjugated anti-EpCAM (CD326) antibodies and anti-PE beads following the manufacturer’s protocol (Miltenyi; Auburn, CA, US).

### Renal Endothelial Cells

Purified primary renal endothelial cells (REC) from C57BL/6 mice were obtained from Cell Biologics and maintained in the Complete Mouse Endothelial Cell Medium (Cell Biologics; Chicago, IL, US). Cells were used between passages 2-6. One million primary REC were combined with 8 μM Control or *Elmo1* On-Target Plus Smart Pool Dharmacon siRNA (Horizon Discovery Biosciences; Cambridge, UK) and transfected using the Human Microvascular Endothelial Cell-Lung Nucleofector Kit (Lonza; Basel, BS) and the Nucleofector 2b device (Lonza), following manufacturer’s instructions. Following transfection, cells were seeded at 20 x 10^4^ cells per well in 24-well tissue culture treated plates. Knockdown efficiency and efferocytosis were analyzed 72 hours post transfection.

### Isolation of renal macrophages

Seventy-two hours after induction of acute kidney injury (AKI) with cisplatin as described above, mouse kidneys were harvested and mechanically dissociated. Tissue was digested in collagenase type I (1.5 mg/mL; Sigma) for 60 min at 37 °C. The digested kidneys were then passed through a sterile 70-μm nylon cell strainer (Corning; Corning, NY, USA) and collected in 50 mL tubes. Cell suspensions were centrifuged at 2000rpm for 10 min, and red blood cells were lysed using Hybri-Max Red Blood Cell Lysing Buffer (Sigma; St. Louis, MO, USA) according to the manufacturer’s instructions. After washing, cells were resuspended in PBS 1X. A discontinuous Percoll gradient (30% and 70%; Cytivia, Marlborough, MA, USA) was prepared, and the cell suspension was carefully layered on top. Gradients were centrifuged at 500g for 30 min at room temperature without brake. Cells at the interphase were collected, washed in PBS 1X, and resuspended in complete RPMI medium supplemented with 10% FBS. To enrich for macrophages, cells were plated and allowed to adhere for 1 hours at 37 °C in a humidified incubator. Non-adherent cells were removed by gentle washing with PBS, and adherent cells were used for efferocytosis assay.

### Efferocytosis

Apoptosis was induced in Jurkat cells by UV-C irradiation (150 mJ/cm², UV Crosslinker, Fisher Scientific) followed by 2 hours of incubation, and in thymocytes by two cycles of UV-C. RTECs were also used as apoptotic targets after cisplatin treatment (75 μM, 48 hours). Apoptosis was confirmed by Annexin V (BD Bioscience #556419) and 7AAD (Thermo Fisher) staining according to the manufacturer’s instructions and analyzed on Attune Nxt flow cytometer (Thermo Fisher) with FCS Express 7 (De Novo Software; Pasadena, CA, US) or FlowJo v10 (BD Bioscience; San Jose, CA, US) software. Apoptotic cells were labeled with 1 μM CypHer5E (GE Healthcare; Chicago, IL, US) and co-incubated with phagocytes—RTECs, RECs, or macrophages—at a 10:1 (Target:Phagocyte) ratio. Co-incubation lasted 2 hours for Jurkat and cisplatin-treated RTEC targets, and 30 minutes for thymocytes with macrophages. Efferocytosis was quantified by flow cytometry and analyzed with FCS Express 7 or FlowJo v10 software.

### Immunoblotting

Proteins from RTECs or RECs were extracted with RIPA lysis buffer with added protease inhibitors cocktail (Roche; Basel, Switzerland). Proteins were separated using SDS-PAGE and transferred onto polyvinylidene fluoride (PVDF) membranes using the Trans-Blot Turbo transfer system (Bio-Rad). The PVDF membranes were blocked with 5% skim milk for 2 h at room temperature and washed three times with Tris-buffered saline containing 0.05% tween-20. The PVDF membranes were incubated with an in-house made anti-ELMO1 rabbit polyclonal antibody (1:1000)(65), anti-beta-Actin-HRP (1:10,000, Sigma #A3854), or anti-ERK1/2 (1:1000, Cell Signaling #9102; Danvers, MA, US), followed by incubation with anti-rabbit secondary antibody (1:10,000, Jackson Immunoresearch # 313-035-003). Blots were exposed using the Western Lightning Plus ECL kit (Perkin–Elmer; Shelton, CT, US) on the ChemiDoc Touch imaging system and analyzed using ImageLab (Bio-Rad).

### ELISA

For protein isolation, the kidney tissue was snap frozen in liquid nitrogen. Tissue extracts were prepared by mechanical dissociation in PBS with added protease inhibitors cocktail (Roche). Kidney tissue extracts and serum were used in a sandwich ELISA assay to determine the level of IL-6 (Thermo Fisher), TNF-α (R&D Systems; Minneapolis, MN, US), neutrophil elastase (R&D Systems) and myeloperoxidase (Thermo Fisher), following manufacturer instructions.

### Strategy for Single-Cell/Nuclei RNA-seq Data Analysis

Single-cell transcriptomic analyses of *ELMO1/Elmo1* expression were performed using publicly available human and mouse kidney datasets. Human single-cell RNA-sequencing data were obtained from the Kidney Precision Medicine Project (KPMP Consortium Dataset)(40). Mouse data were retrieved from(41) (GEO accession GSE139107), which profiled kidneys following ischemia–reperfusion injury (IRI) at control, 4 hours, 12 hours, 2 days, 14 days, and 6 weeks post-injury. All datasets were analyzed in R (version 4.4) using the Seurat(66), SeuratObject, tidyverse, ggplot2, and gridExtra packages. Raw digital gene-expression (DGE) matrices from the mouse study were imported as individual Seurat objects, annotated using accompanying metadata, and subjected to quality-control filtering. Datasets were normalized by log transformation, highly variable genes were identified, and integration anchors were calculated to correct for batch and time-point effects, generating a unified Seurat object. For the human KPMP dataset, the processed .h5Seurat object was imported directly, and cell annotations were retained as provided by the KPMP consortium. Disease groups were categorized as Control (Reference), acute kidney injury (AKI), and chronic kidney disease (CKD). For both species, *ELMO1/Elmo1* expression was evaluated across annotated cell types and conditions.

### GWAS and linkage analysis

We searched publicly available genome-wide association studies of human disease for single nucleotide polymorphisms (SNP) by gene interactions(67). The list of curated databases that were used is summarized in ***Supplementary Table 1***. Significance of a given SNP was determined in the original study. SNP by phenotype interactions were determined using the GWASdb SNP-phenotype association database. The data are plotted using a standardized *p* value as reported(68). For determination of experimental linkage, a search of curated databases such as the DISEASES experimental gene-disease association database was performed. Significance of the experimental linkage association was determined via a previously established aggregated score(69–73).

### Statistical analysis

Statistical significance was determined using GraphPad Prism 9 using an unpaired Student’s two-tailed t-test. The variance was similar between groups. No inclusion/exclusion criteria were pre-established. A p-value of <0.05 (indicated by one asterisk), <0.01 (indicated by two asterisks), <0.001 (indicated by three asterisks), and <0.0001 (indicated by four asterisks) was considered significant.

### Illustration

For the preparation of figures, we have modified illustrations available through www.motifolio.com or used BioRender.

## Acknowledgements

The authors thank members of the Arandjelovic laboratory and the University of Virginia Center for Immunity, Inflammation, and Regenerative Medicine (CIIR) for discussions, and Alban Gaultier and Lisa Haney for critical reading of the manuscript. This work is supported by funding to S.A. from the NIH R01AI158596 and the University of Virginia.

## Author Contributions

Conceptualization, B.B., M.C. and S.A.; Methodology, B.B., M.C., V.S. and S.A.; Software, V.S.; Investigation, B.B, M.C., V.S., P.M., I.H., S.Z., J.G., K.S., L.S., M.O., R.S. and S.A.; Data Curation, I.H. and V.S.; Writing, B.B. and S.A.; Funding Acquisition, S.A. All authors read and approved the final paper.

## Ethics Statement

This study did not utilize human participants that would require ethical approval. All animal procedures were approved by and performed according to the guidelines of the Institutional Animal Care and Use Committee (IACUC) at the University of Virginia under the protocol #4320.

## Data Availability Statement

The data that support the findings from this study, including the R code used for bioinformatics analysis, are available from the corresponding authors upon reasonable request.

