## Supplementary Information for "ELMO1 dependent efferocytosis protects from nephrotoxin induced acute kidney injury"

**Supplementary Figure 1.** ***Elmo1*$^{-/-}$ mice have normal kidney function at baseline.** (**A**) Body weight (g) and kidney weight (g) of healthy *Elmo1*$^{+/+}$ (n=32, 4) and *Elmo1*$^{-/-}$ mice (n=22, 4). (**B**) Serum creatinine and blood urea nitrogen (BUN) levels in healthy *Elmo1*$^{+/+}$ (n=5) and *Elmo1*$^{-/-}$ (n=4) mice. (**C**) Kidney morphology assessed by hematoxylin-eosin (H&E) staining in healthy mice. Representative images are shown. Scale bars indicate 1 mm. Data are shown as mean ± SD; p values were determined by unpaired Student's *t*-test; ns, not significant. Each symbol indicates an individual animal.

**Supplementary Figure 2.** ***Elmo1***$^{-/-}$ **mice have reduced inflammatory response after IRI-AKI.** (**A**) Kidney pathology at 24 h post IRI-AKI, assessed by hematoxylin-eosin (H&E) staining. Representative images are shown alongside quantification of kidney pathology scores in *Elmo1*$^{+/+}$ (n=4) and *Elmo1*$^{-/-}$ (n=3) mice. Scale bars indicate 1 mm. Data are from one experiment. (**B**) Relative gene expression of inflammatory markers (*Il-6, Tnfα, Il-10,* and *Il-18*) in kidneys from *Elmo1*$^{+/+}$ (n=11-20) and *Elmo1*$^{-/-}$ (n=7-21) mice following IRI-AKI, normalized to *Hprt1*. Data shown are pooled from more than 3 independent experiments. Each symbol indicates an individual animal. (**C**) Protein levels of IL-6 and TNFα in kidney tissue from *Elmo1*$^{+/+}$ (n=4) and *Elmo1*$^{-/-}$ (n=3) mice on Day 1 after IRI-AKI. Data shown are from one experiment. Each symbol indicated an individual animal. (**D**) Protein levels of IL-6 and TNFα in serum from *Elmo1*$^{+/+}$ (n=4-9) and *Elmo1*$^{-/-}$ (n=3-6) mice following IRI-AKI. Data shown are pooled from two experiments. Each symbol indicated an individual animal. (**E**) Protein levels of neutrophil markers elastase and myeloperoxidase (MPO) in serum from *Elmo1*$^{+/+}$ (n=9) and *Elmo1*$^{-/-}$ (n=6) mice following IRI-AKI. Data shown are pooled from two experiments. Each symbol indicated an individual animal. (**F**) Protein levels of neutrophil markers elastase and MPO in kidney tissue from *Elmo1*$^{+/+}$ (n=4) and *Elmo1*$^{-/-}$ (n=3) mice following IRI-AKI. Data shown are from one experiment. Each symbol indicated an individual animal. (**G**) Representative images of neutrophil infiltration in kidney sections, assessed by Ly6G staining (red), from *Elmo1*$^{+/+}$ (n=3) and *Elmo1*$^{-/-}$ (n=3) mice following IRI-AKI. Nuclei were stained with Hoechst (blue). Data is presented as percentage of total tissue area. Scale bar indicates 1 mm. Data are from one experiment. Data are shown as mean ± SD; p values were determined by unpaired Student's *t*-test; $*p < 0.05$.

**Supplementary Figure 3.** ***Elmo1*$^{-/-}$ kidney inflammatory gene expression in cisplatin-AKI.** Relative gene expression of inflammatory markers (*Il-6, Tnfα and Il-10*) in the kidneys from *Elmo1*$^{+/+}$ (n=24), *Elmo1*$^{-/-}$ (n=18), and control$^{+/+}$ (n=5) mice following cisplatin-AKI, normalized to *Hprt1*. Data are pooled from 3 independent experiments. Each symbol indicated an individual animal.

**Supplementary Figure 4. Deletion of *Elmo1* does not increase cisplatin-induced cell death in renal tubular epithelial cells (RTEC).** (**A**) Flow cytometry gating strategy and purity assessment of cultured primary renal tubular epithelial cells (RTEC). (**B**) Quantification of the percentage of $EpCAM^+$ cells in primary RTEC cultures from $Elmo1^{+/+}$ (n=11) and $Elmo1^{-/-}$ (n=11) mice. Data are pooled from 3 independent experiments. Data are shown as mean ± SEM; p values were determined by unpaired Student's *t*-test. ns, not significant. (**C**) Immunofluorescence staining of cultured primary RTEC for ZO-1 (green) to visualize tight junctions. Hoechst (blue) was used to stain the nuclei. Scale bar, 100 µm. Representative images are shown from one experiment with primary RTEC from $Elmo1^{+/+}$ (n=3) and 3 $Elmo1^{-/-}$ (n=3) mice. (**D**) Immunoblot of protein expression of ELMO1 in cultured RTEC from $Elmo1^{+/+}$ (n=4) and $Elmo1^{-/-}$ (n=3) mice. Actin was used for protein loading control. Each lane represents primary cells established from a separate animal. (**E**) Representative flow cytometry plots of Annexin V and 7AAD stained RTEC from $Elmo1^{+/+}$ (n=3) and $Elmo1^{-/-}$ (n=3) mice after 48 hours of treatment with 125 µM cisplatin. (**F**) Dose-response of cisplatin-induced apoptosis ($AnnexinV^+7AAD^-$) over 48 h in RTEC, with concentrations of 0-125 µM. Graphs show percentage of apoptotic cells. Data are pooled from 2 independent experiments.

**Supplementary Figure 5. Cleaved caspase-3 staining does not overlap with glomeruli in cisplatin-AKI.** Representative images of kidney sections from cisplatin-AKI mice stained for cleaved caspase-3. Scale bars indicate 1 mm. Arrows point to glomeruli.

### Supplementary Figure 1. *Elmo1$^{-/-}$* mice have normal kidney function at baseline.

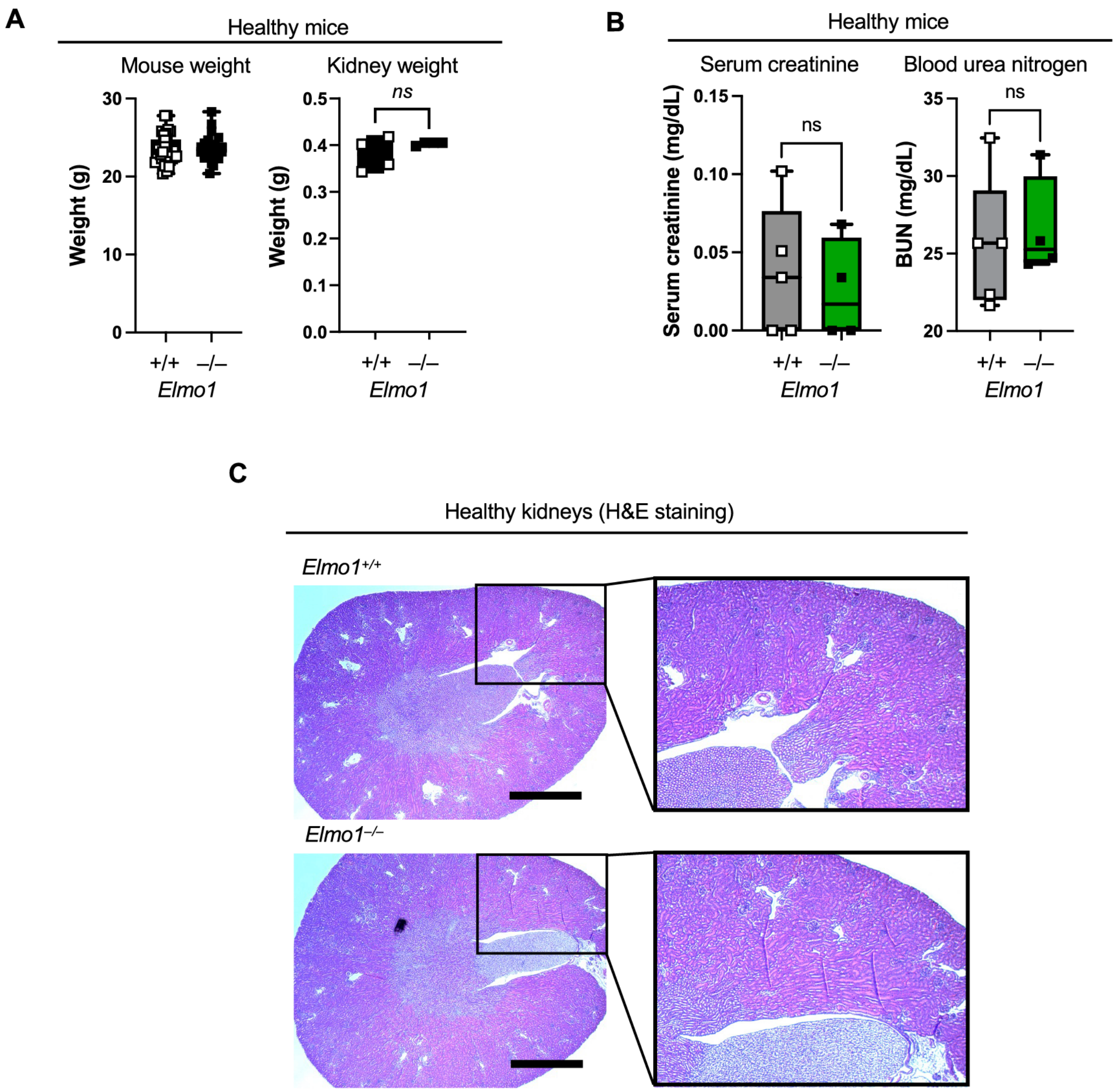

### Supplementary Figure 2. *Elmo1*$^{-/-}$ mice have reduced inflammatory response after IRI-AKI.

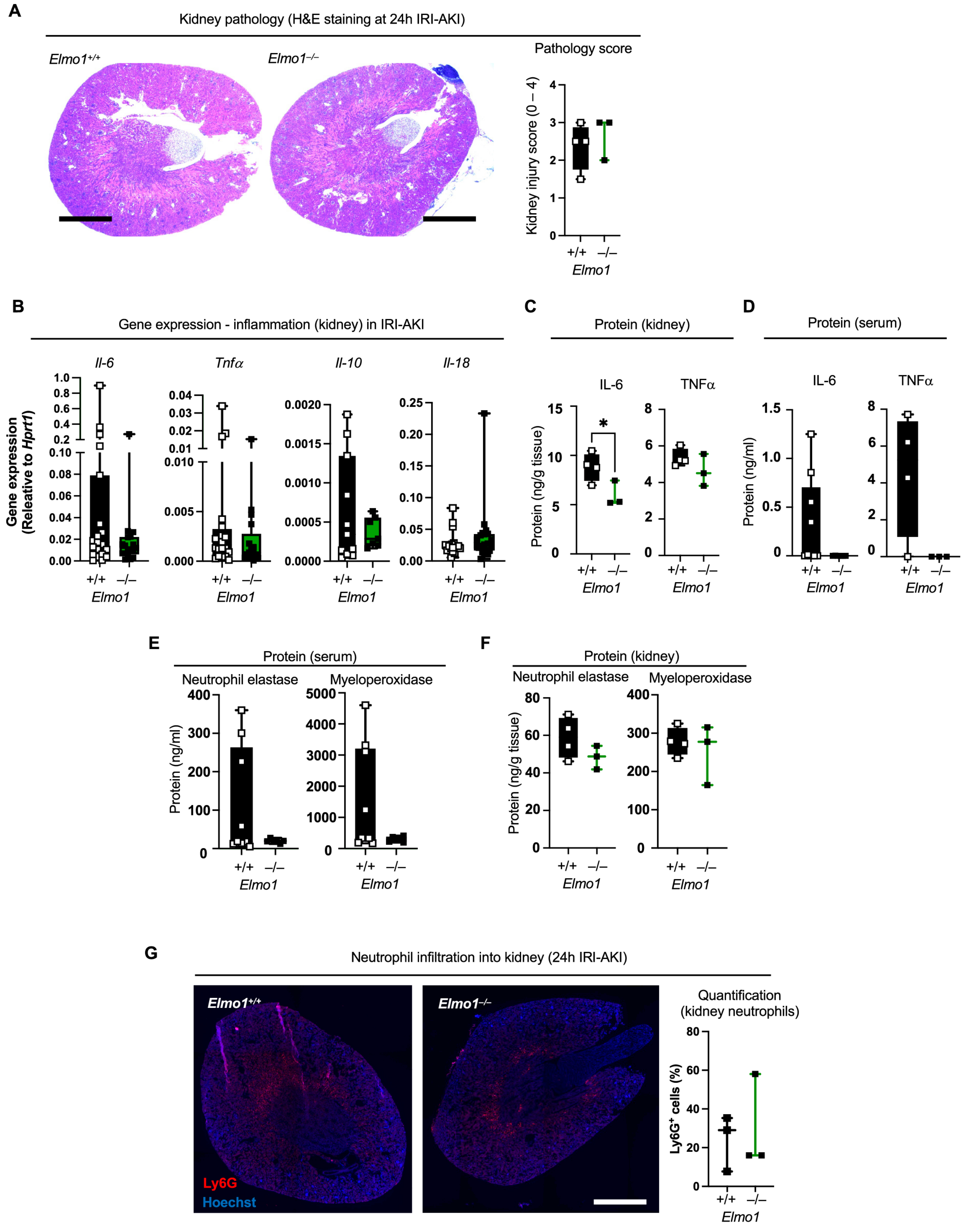

### Supplementary Figure 3. *Elmo1*$^{-/-}$ kidney inflammatory gene expression in cisplatin-AKI.

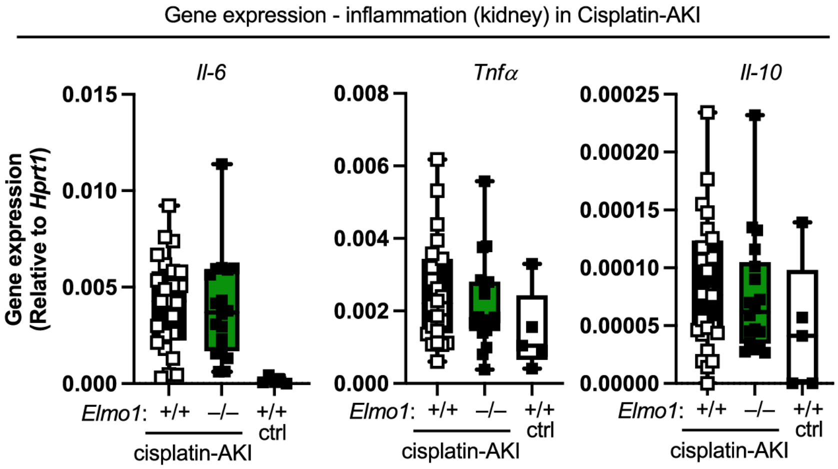

### Supplementary Figure 4. Deletion of *Elmo1* does not increase cisplatin-induced cell death in renal tubular epithelial cells (RTEC).

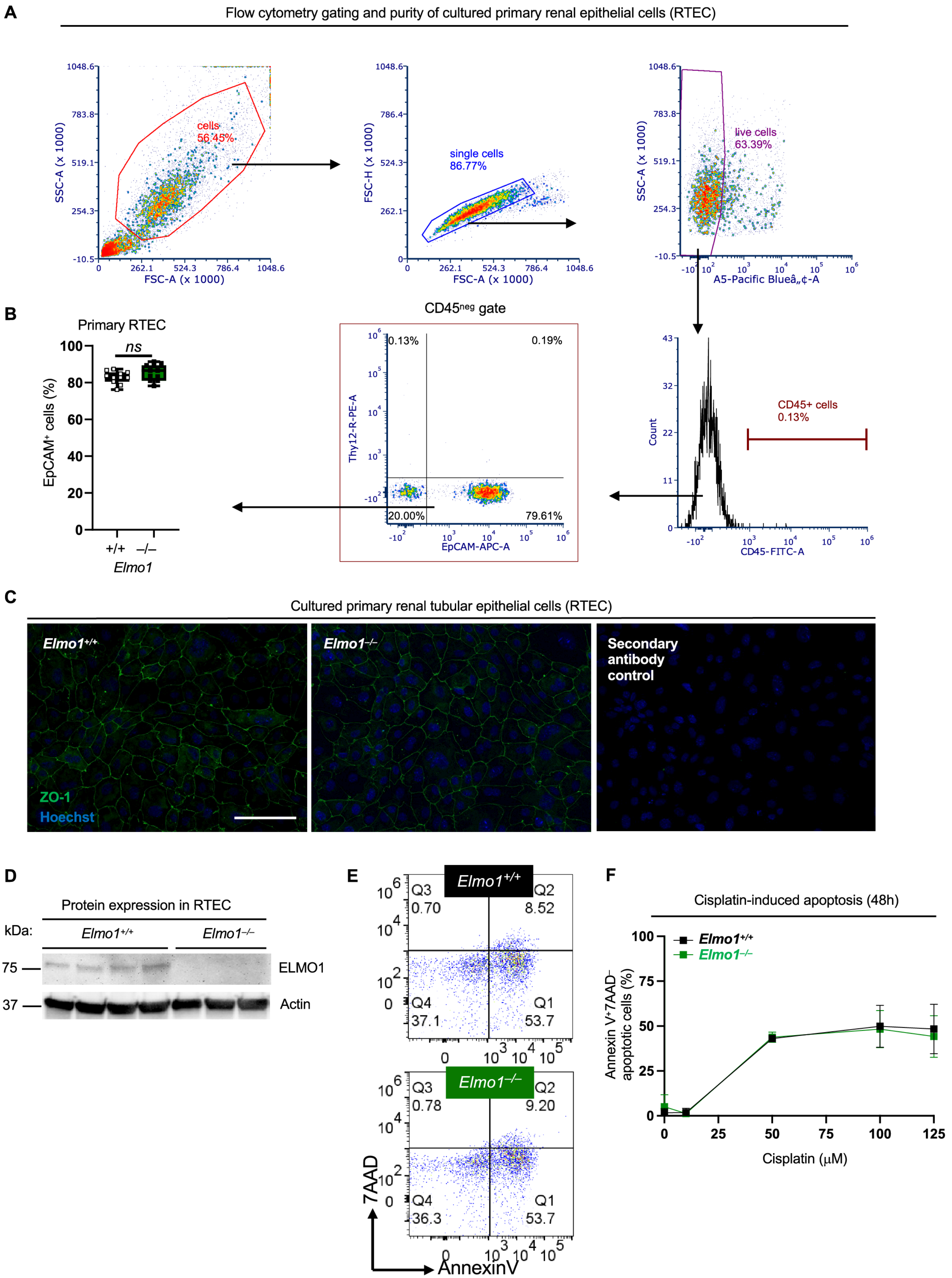

### Supplementary Figure 5. Cleaved caspase-3 staining does not overlap with glomeruli in cisplatin-AKI.

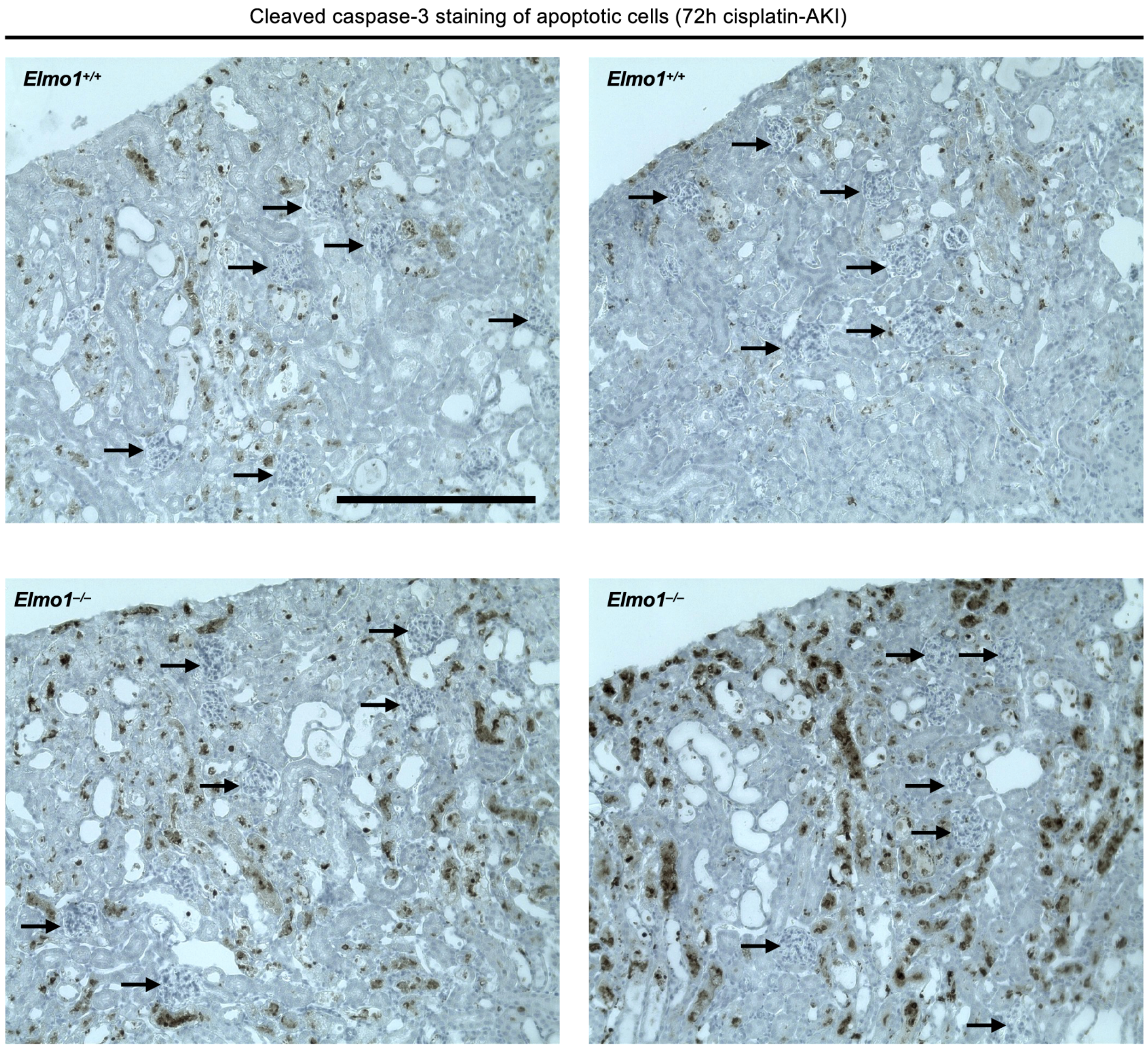

***Supplemetary Table 1.*** List of Databases Used.

| **Database Name** | **DOI of Originial Paper** |
|---|---|
| BioGPS Cell Line Gene Expression Profiles | 10.1073/pnas.0400782101 |
| | 10.1073/pnas.012025199 |
| | 10.1093/nar/gks1114 |
| CTD Gene-Disease Associations | 10.1093/nar/gku935 |
| | 10.1093/nar/gkn580 |
| DISEASES Experimental Gene-Disease Assocation Evidence Scores | 10.1016/j.ymeth.2014.11.020 |
| DISEASES Text-mining Gene-Disease Assocation Evidence Scores | 10.1016/j.ymeth.2014.11.020 |
| GAD Gene-Disease Associations | 10.1038/ng0504-431 |
| GeneRIF Biological Term Annotations | PMC1480312 |
| GTEx Tissue Gene Expression Profiles | 10.1126/science.1262110 |
| | 10.1038/ng.2653 |
| GWAS Catalog SNP-Phenotype Associations | 10.1093/nar/gkt1229 |
| GWASdb SNP-Disease Associations | 10.1093/nar/gkr1182 |
| GWASdb SNP-Phenotype Associations | 10.1093/nar/gkr1182 |
| HuGE Navigator Gene-Phenotype Associations | 10.1038/ng0208-124 |
| TISSUES Curated Tissue Protein Expression Evidence Scores | 10.7717/peerj.1054 |

| **Database Tool** | **DOI** |
|---|---|
| Harmonizome | 10.1093/database/baw100 |
